# The RNA binding protein Pub1 inhibits TORC1 activity in *Saccharomyces cerevisiae*

**DOI:** 10.1101/2024.08.22.608743

**Authors:** Helena Orozco, Emilia Matallana, Mikael Molin, Agustín Aranda

## Abstract

Pub1 is an RNA binding protein with multiple functions in gene expression, including the control of translation and mRNA stability. It also acts as a prion-like protein given its ability to self-aggregate, and it plays a role in the formation of mRNA-protein assemblies such as stress granules. This study identifies an unexpected relationship between Pub1 and the nutrient signalling complex TORC1. Whereas Pub1-deficiency increased cellular resistance to TORC1 inhibition by rapamycin, *PUB1* overexpression impaired growth, particularly when nutrients were scarce. This growth defect was suppressed by *TOR1* kinase deletion, which indicates that Pub1 functions are channelled through the TORC1 complex. Moreover, Pub1 antagonised TORC1-regulated processes, such as autophagy and the phosphorylation of ribosomal protein Rps6. We found that Pub1 was required for the aggregation of Kog1, a component of the TORC1 complex upon starvation, whereas its overexpression caused Kog1 foci formation in non-stressed growing cells. All these results suggest a negative role of Pub1 in regulating TORC1 activity based on its ability to promote the formation of TORC1-ribonucleoprotein aggregates.

## Introduction

Nutrient-sensing pathways coordinate all cellular physiological processes in response to nutrient supply (1). One of the main pathways of this kind in eukaryotic cells has been proposed to involve the TORC1 complex. TOR proteins are highly conserved throughout eukaryotes and share their homology with other phosphatidyl inositol kinases (2). The budding yeast *Saccharomyces cerevisiae* has two different TOR proteins: Tor1, which is a component of the rapamycin-sensitive TORC1 complex, and the essential Tor2, which can take part in both TORC1 and the rapamycin-insensitive TORC2 complex. TORC1 is thought to respond to certain nutrients and thereby regulates a diverse set of cellular processes, including growth, aging, metabolism, autophagy and stress response (2, 3). TORC1 integrates different stimuli and its outcome is different if it is inhibited chemically by rapamycin or inactivated by nutritional deficiencies such as nitrogen starvation, nitrogen limitation or leucine starvation (4). Inhibition of TORC1 by rapamycin triggers many processes related to the recycling of cellular molecules, including a process called macroautophagy that involves degradation in the vacuoles of large portions of the cytoplasm in structures called autophagosomes (5). This is one of the processes involved in antagonising aging. Inhibition of TORC1 signalling by amino acid depletion, treatment with drugs that inhibit TORC1 activity such as rapamycin or the deletion of components of the signalling pathway lead to enhanced chronological life span (CLS) (6), the ability of cells to remain viable in a non-dividing state, such as the stationary phase (7). Reduction of TORC1 signalling improves cellular resistance to heat and oxidative stress and induces the nuclear localisation of stress response transcription factor Msn2. Stress also controls TORC1, in many cases by controlling its aggregation. It has been described that upon heat shock, the TORC1 complex is sequestered transiently in stress granules (SG) (8). Similarly upon glucose depletion, one of the components of TOR complexes, Raptor protein Kog1, aggregates in foci called Kog1-bodies and causes the down-regulation of TORC1 activity (9). This is mediated by another nutrient-sensing kinase, AMPK Snf1, that is involved in glucose repression and stress signalling (1). Recently it has been described up to 13 activators of TORC1 aggregation, and seven of them work through Gtr1/2 GTPases, a well-known TORC1 regulator (10). TORC1 complexes also oligomerise under starvation into a hollow helical assembly called TOROID (11). Recently it has been described that yeast ataxin-2 Pbp1 (a component of SG) is able to form an intracellular condensate under respiratory conditions that inhibits TORC1, inducing autophagy in a prototrophic strain (12).

Stress granules are ribonucleoprotein (RNP) structures that form in response to a variety of environmental conditions (13, 14). They are considered places of transient sequestration of proteins involved in translation and mRNA, whose dissolution allows translation to be resumed once the environmental insult disappears. Reflecting this, the polyA binding protein Pab1 is a hallmark component of such granules. Pub1 is an mRNA-binding protein (RBP) and an SG component required for SG formation under glucose deprivation, but not azide stress (15). It has a prion-like ability to self-aggregate, which enables the formation of a self-propagation aggregate with translation termination protein Sup35 (16). Pub1 has the ability to act as a translational inhibitor in a carbon source-dependent way (17). It also modulates the activity and localisation of glycerol synthesis protein Gpd1 (18). The many functions of Pub1 may explain its impact on cell longevity under different environmental conditions (19).

This work analysed the relationship between the mRNA binding protein Pub1 and the TORC1 complex in different growth conditions, including starvation and chronological ageing. The analysis of growth, rapamycin resistance, autophagy and the phosphorylation of Rps6 in cells lacking or overexpressing *PUB1* suggest that Pub1 inhibits TORC1. Furthermore, the effects of *PUB1* on CLS depend on the interaction of both. These phenotypes may be related to an aberrant sequestration of the TORC1 complex or TORC1 components such as Kog1 due to Pub1 activity.

## Results

### RBPs are autophagy activators

Our previous works studied the role of SG component Pub1 in stress tolerance, aging and glycerol production, and suggested an interconnection between Pub1 and Tor1 in the control of chronological longevity (18, 19). Autophagy is a well-known quality control mechanism which correlates with longevity and that is repressed by the TORC1 nutrient-sensing pathway. In order to analyse the potential impact of SG components on autophagy, we deleted the genes encoding for its components Pub1, Ngr1 and Pbp1 in the prototrophic strain C9, a wine yeast-derived strain previously used to study *PUB1* under different environmental conditions (18, 19). To follow autophagy, we measured the degradation and release of GFP of the glycolytic enzyme Pgk1-GFP fusion protein by Western blot (20). We tested the deletion mutants of *PUB1*, *NGR1* and *PBP1* together with a *TOR1* deletion strain under two conditions that induce autophagy: presence of rapamycin and nitrogen starvation (Figure 1). Figure 1A shows that both mutants *pub1*Δ and *ngr1*Δ failed to cleave the GFP signal in response to rapamycin. Therefore, they play a positive role in promoting autophagy under this condition. However, the *pbp1*Δ mutant shows normal Pgk1-GFP processing, which indicates that it does not contribute to autophagy under these conditions. As expected, the *tor1*Δ mutant that exhibited decreased TORC1 activity displayed a larger amount of cleaved GFP, which indicates increased autophagy. Behaviour was similar upon nitrogen starvation (Figure 1B), but in this case, *ngr1*Δ and *pbp1*Δ mutants instead displayed a diminished Pgk1-GFP cleavage, while almost normal autophagy appeared to take place in the *pub1*Δ mutant judging by the reduced levels of cleaved GFP. The fact that *TOR1* deletion gives almost normal levels of autophagy indicate that this condition is quite different and may trigger alternative pathways that promote autophagy in the long run. Therefore, RBP components of SG play a positive role in autophagy under different environmental stress conditions, but their contributions to the process may vary, in a similar way that these different conditions inhibit TORC1 in a different way. Overall it is noteworthy that RBP components seem to play the opposite role to that of the TORC1 complex in this process.

**Figure 1.**
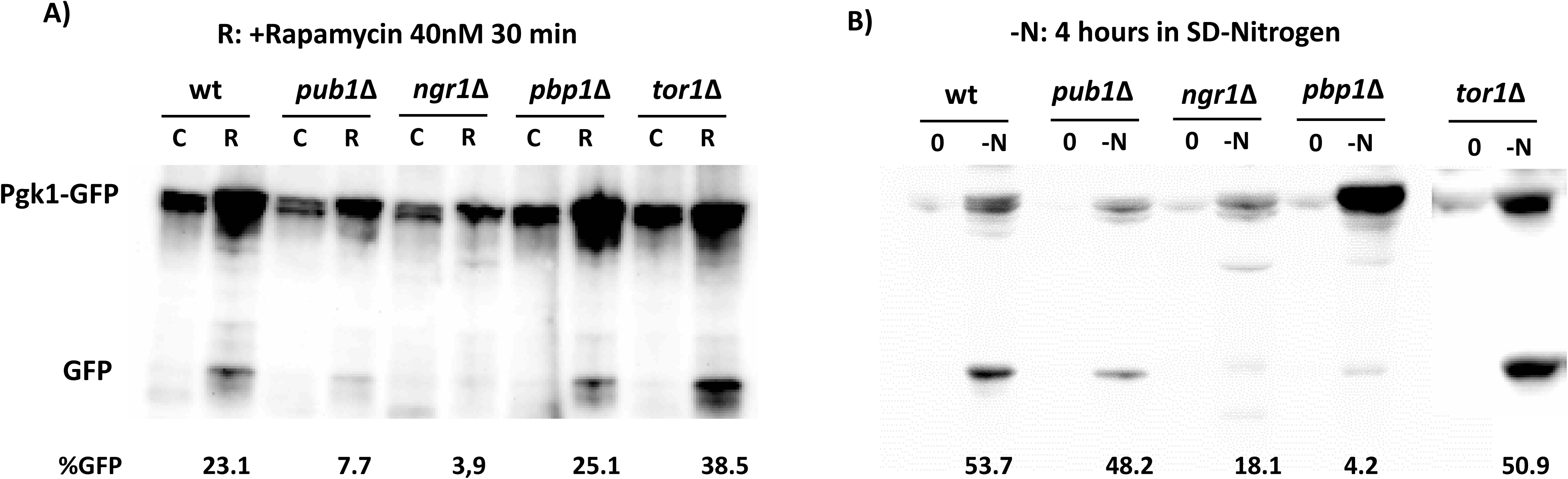
RNA binding proteins of Stress Granules play a positive role in autophagy. A) The Western blot of the wild-type C9 strain and the deletion mutants of genes *PUB1*, *NGR1*, *RBP1* and *TOR1* also expressing a GFP tagged version of Pgk1. Cells were grown exponentially in SD medium (SD) and treated with 40 nM rapamycin for 30 minutes (R) or were not treated as the control (C). The fusion protein and GFP fragment that marks degradation by autophagy are indicated. The percentage of cleaved GFP referred to the total GFP+Pgk1-GFP is indicated for each mutant. B) The same strains as in panel A) were incubated in a SD medium without nitrogen (-N) for 4 h to trigger autophagy.

### Pub1 is functionally linked to the TORC1 complex

In order to further analyse the potential link between the RBPs belonging to SG to the TORC1 complex, cellular resistance against the TORC1 inhibitor rapamycin of strains deleted for *PUB1*, *NGR1* and *PBP1* was tested. A spot test analysis was performed on plates of rich medium containing 100 nM of rapamycin (Figure 2A). As a control, a *TOR1* deletion strain was included, which displayed the expected hypersensitivity caused by reducing TORC1 complex activity. Interestingly, the *pub1*Δ strain displayed a moderately increased resistance to rapamycin, which may suggest that Pub1 negatively regulates TORC1 activity. In contrast, *pbp1*Δ and *ngr1*Δ mutants grew as the wild type and therefore we focused on Pub1 from this point onwards. To substantiate the apparent rapamycin hyperresistance by a different approach, we measured growth in the rich medium YPD supplemented with a lower amount of rapamycin, 1 nM, of both deletion mutants *TOR1* and *PUB1*, and of double mutant *pub1*Δ*tor1*Δ (Figure 2B). In the absence of rapamycin, all strains display a similar growth profile and reached the same final density. Therefore, none of these mutations impaired growth under optimal conditions. However, even with this low level of rapamycin, the *TOR1* deletion strain displayed a dramatic growth defect. The strain lacking Pub1 exhibited a slight delay compared to the parental strain, suggesting that Pub1 function stimulates growth in the presence of rapamycin. However, this strain reached a higher cell density, indicating that the Pub1 function would be positive upon stationary phase or when nutrients become limiting and the TORC1 complex is less active. This suggests that Pub1 is inhibiting TORC1 at the entry into stationary phase. The double mutant had a similar profile as the *TOR1* deletion strain, which suggests that Pub1 may function upstream of TORC1 and modulate its activity.

**Figure 2.**
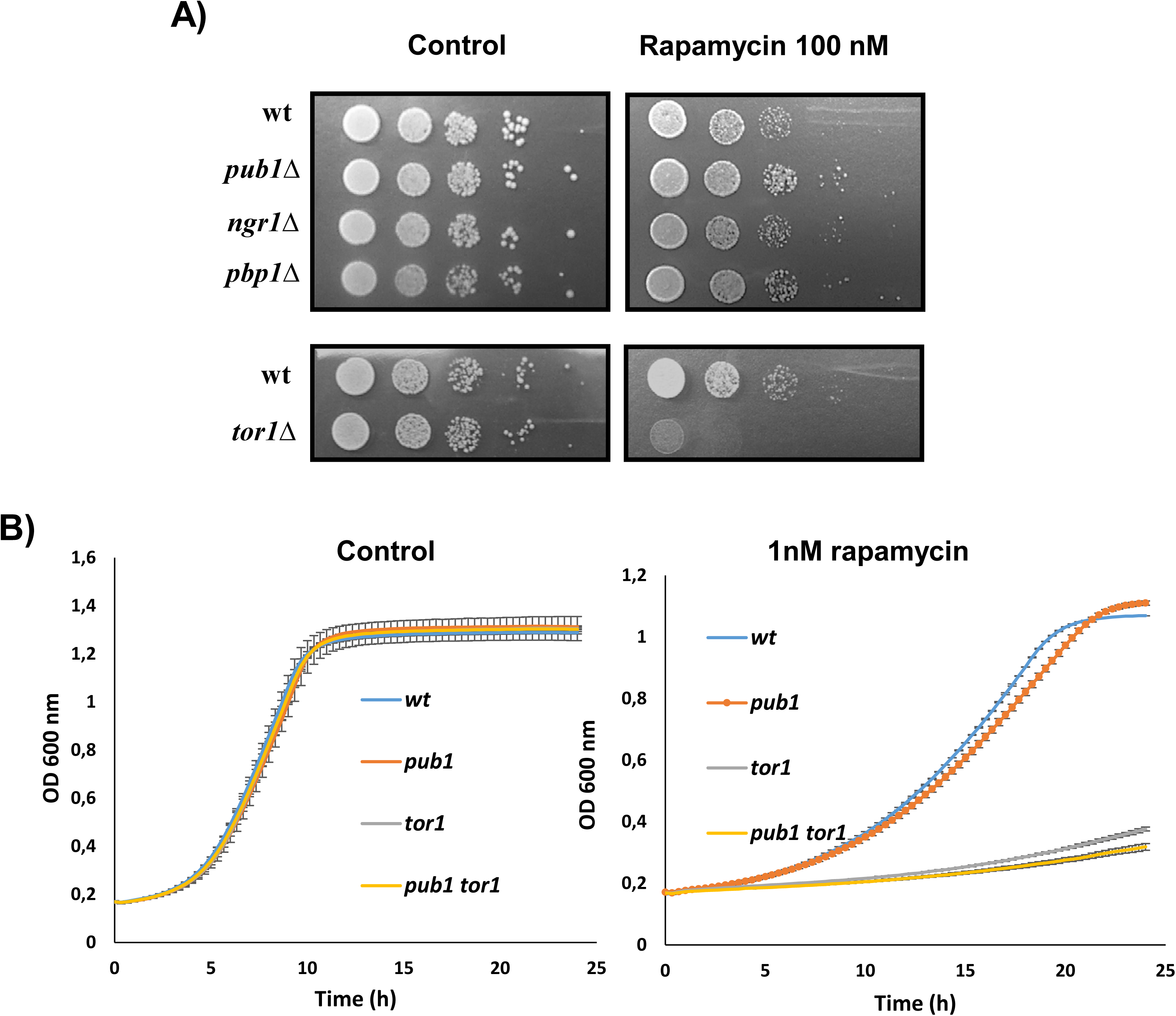
Pub1 inhibits TORC1 activity. A) The spot test analysis of the serial 10-fold dilutions of the wild-type and *pub1*Δ, *ngr1*Δ, *pbp1*Δ and *tor1*Δ mutants on YPD medium with or without 100 nM rapamycin. B) Growth curves on YPD and YPD + 1nM rapamycin of the *pub1*Δ, *tor1*Δ and *pub1*Δ*tor1*Δ mutants measured by a Varioskan multi-well plate reader with shaking and incubation at 30°C. Experiments were done in triplicate. The mean and standard deviation are provided.

### *PUB1* overexpression decreases growth and longevity

In order to better understand the role of Pub1 and its interaction with TORC1, we overexpressed the *PUB1* gene by replacing its promoter with the methionine-controlled *MET17* promoter, a modification that allows its overexpression in a controlled manner. This approach has previously allowed us to overexpress other genes of interest (19, 21). We performed this overexpression also in a *TOR1* deletion mutant to further characterise the interaction between these two proteins. Cells were grown in a synthetic complete SC medium lacking methionine where, according to the qPCR analysis, *PUB1* was overexpressed 3-fold (Supplementary Figure S1). Cell viability was measured over time to obtain a growth curve (Figure 3A). This is the medium for standard CLS measurements (22); thus a CLS plot can be obtained if day 3 is taken as 100% viability (Figure 3B). *TOR1* deletion does not cause any defect in growth (Figure 3A). Under these conditions, *TOR2* may be able to constitute enough TORC1 to promote growth. *PUB1* overexpression leads to reduced cell growth and the culture achieved a lower final cell density, which indicates that its overexpression alters cellular growth. However, the combination of these two genetic modifications led to a strain that grew almost as well as the reference strain and the *tor1*Δ strain. Once again, this indicates that Pub1 may channel some of its functions through the TORC1 complex. In the context of ageing, *TOR1* deletion extended life span (Figure 3B), as shown in many cases in this and other genetic backgrounds (6, 23). *PUB1* overexpression caused a drop in the life span, which indicates that it is impacting processes in the stationary phase. The *tor1*Δ *MET17-PUB1* double mutant gave an intermediary CLS pattern, similarly to the reference strain. This implies again that Pub1 antagonizes TORC1 activity.

**Figure 3.**
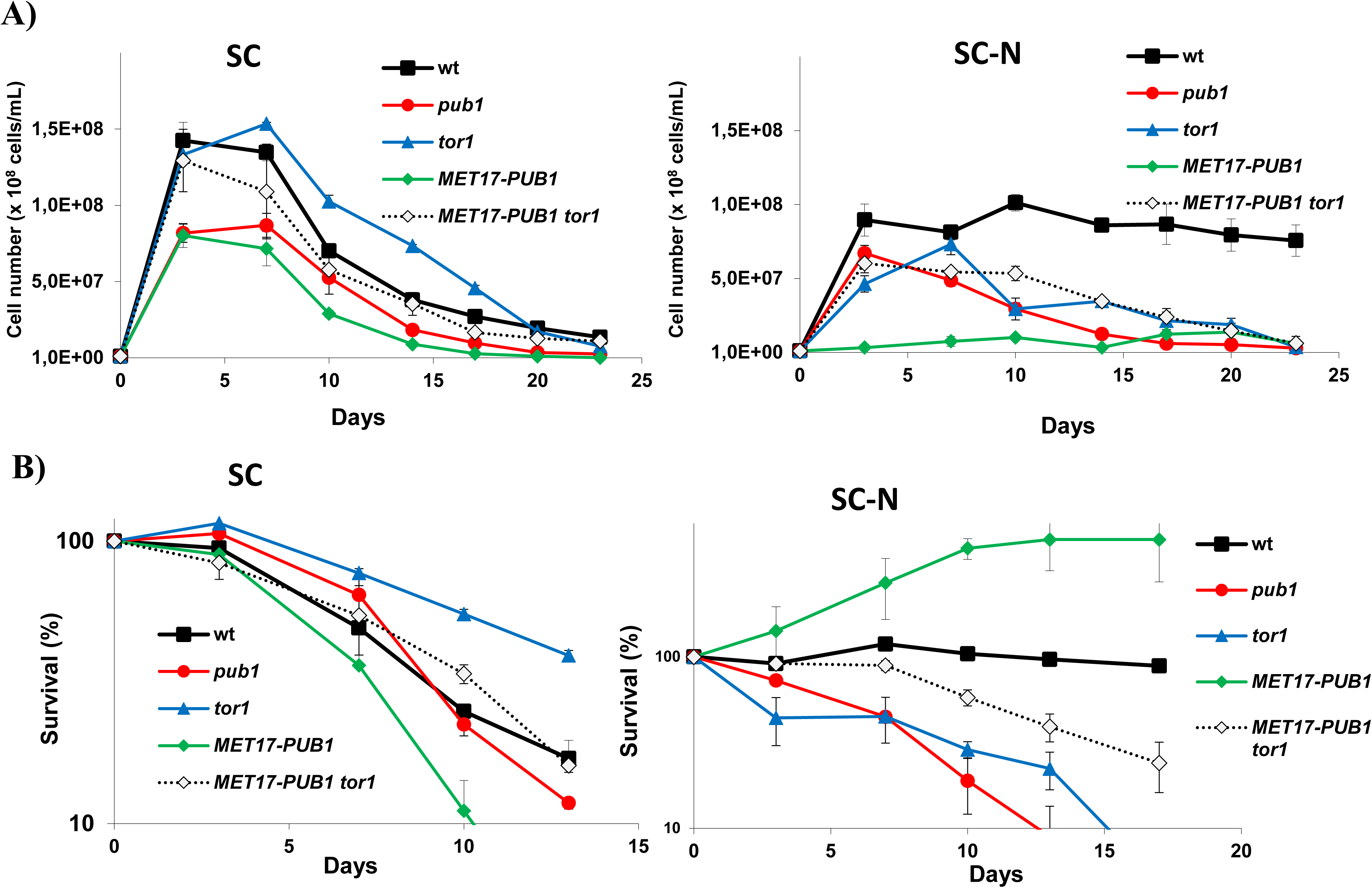
Cells overexpressing *PUB1* display a growth defect. A) Viability count of the growth cultures on SC minus methionine of the wild-type, *tor1*Δ, *MET17-PUB1* and *MET17-PUB1 tor1*Δ strains. Β) The chronological life span (CLS) plot of panel A considering day 3 to be 100% viability. C) Viability count of the same strains as panel A) in SC with 25-fold less ammonium sulphate and the amino acid drop out mix. D) The CLS plot from the viability count in panel C) when considering day 7 of growth to be 100% viability. Experiments were done in triplicate. The mean and standard deviation are provided.

In previous works, a synthetic medium SC containing 1/25^th^ of amino acids and ammonia was used to create a metabolic situation when nitrogen, and not carbon, was the limiting nutrient (23, 24). Growth was reduced under this condition (Figure 3C) and day 7 was taken as the start of the ageing plot (Figure 3D). In these experiments, *TOR1* deletion delayed growth, probably because nitrogen scarcity may cause TORC1 activity to be poor and *TOR1* deletion aggravates this effect. Upon Pub1 overexpression, cells displayed a dramatically arrested growth, which was rescued by *TOR1* deletion. These findings reinforce the genetic interaction between these two genes. In aging terms, the pattern was similar; the *tor1*Δ strain displayed a shorter life span under these conditions, as observed before (23). The CLS plot of the *MET17-PUB1* strain revealed an abnormal behaviour of this mutant, with a few cells still growing at a slow rate. This situation was reverted in the *tor1*Δ *MET17-PUB1* double mutant by resulting in accelerated ageing compared to the wild type and, thus, displayed a phenotype that was more similar to the *TOR1* deletion strain than to *PUB1* overexpression.

### The poor growth of the cells overexpressing Pub1 is independent of global translation initiation

As Pub1 is present in the 77S translational complex (25) and has been linked to the translational repression of specific transcripts (17), its role in global translation was assayed by a polysome analysis (Table 1, Supplementary Figure S2). Exponentially growing wild-type cells have many ribosomes engaged in translation in the form of polysomes (85.5%). Similar numbers were obtained in cells deletion for *TOR1* (84.95). *PUB1* overexpression did not change the profile, which suggests that its impact on growth was not a consequence of a global arrest in translational initiation. The deletion of *PUB1* slightly increased the percentage of polysomes, which confirms that it may be involved in the repression of the translation of selected transcripts during growth in glucose.

**Table 1.** The polysome analysis of *PUB1* deletion and overexpression. A single experiment was performed for each strain and condition. The percentage of ribosomes present in polysomes is shown for the strains, indicated during exponential growth and after treatment with 200 nM rapamycin for 15 minutes. The ratio between both conditions is shown.

| Strain | Exponential growth | Rapamycin | Ratio |
| --- | --- | --- | --- |
| wt | 85.5 | 66.4 | 0.78 |
| *tor1Δ* | 84.9 | 66.9 | 0.79 |
| *pub1Δ* | 90.7 | 72.9 | 0.80 |
| *MET17-PUB1* | 85.8 | 75.2 | 0.88 |
| *MET17-PUB1 tor1Δ* | 87.7 | 79.6 | 0.91 |

Adding rapamycin for 15 minutes, a condition that decreases ribosome biogenesis (26), lowered the number of polysomes in the wild type by 22.3%. A similar drop was evidenced in the *PUB1* deletion mutant. The reduction in the *tor1*Δ strain was slightly lower (17.7%), probably due to the fact cells had less TORC1 complex to be inhibited by rapamycin. Interestingly, when *PUB1* was overexpressed, translation instead seemed less sensitive to rapamycin (a 12.4% reduction), thus supporting the idea that it has a negative role in regulating TORC1 activity.

### Study of the relationship of Pub1 with different nutrient signalling pathways

Next, in order to confirm the relationship between overexpression and TORC1 activity, the sensitivity of the *PUB1* overexpressing strain to rapamycin was tested, together with the *TOR1* deletion strain and the *tor1*Δ *MET17-PUB1* double mutant (Figure 4). The test was carried out in minimal medium SD plates to achieve *MET17* promoter activation and to test the molecules that inhibit specific steps in amino acid biosynthesis. On this minimal medium, the *MET17-PUB1* overexpression mutant already displayed impaired growth in the control plate, but was rescued when combined with *TOR1* deletion, which also happened in the CLS experiments in liquid media (Figure 3). When a standard concentration of 100 nM rapamycin was tested, all mutant strains displayed strongly impaired growth, which confirmed that these factors played a role in regulating TORC1 activity. Rapamycin was lowered to 5 nM with similar results, and finally to 1 nM when the growth of the wild-type strain was almost normal. Under these conditions, the *TOR1* deletion mutant exhibited moderately increased sensitivity, which became exacerbated in the *tor1*Δ *MET17-PUB1* strain. This reinforces the idea that Pub1 represses TORC1 activity. The *MET17-PUB1* mutant displayed almost negligible growth at all the concentrations.

**Figure 4.**
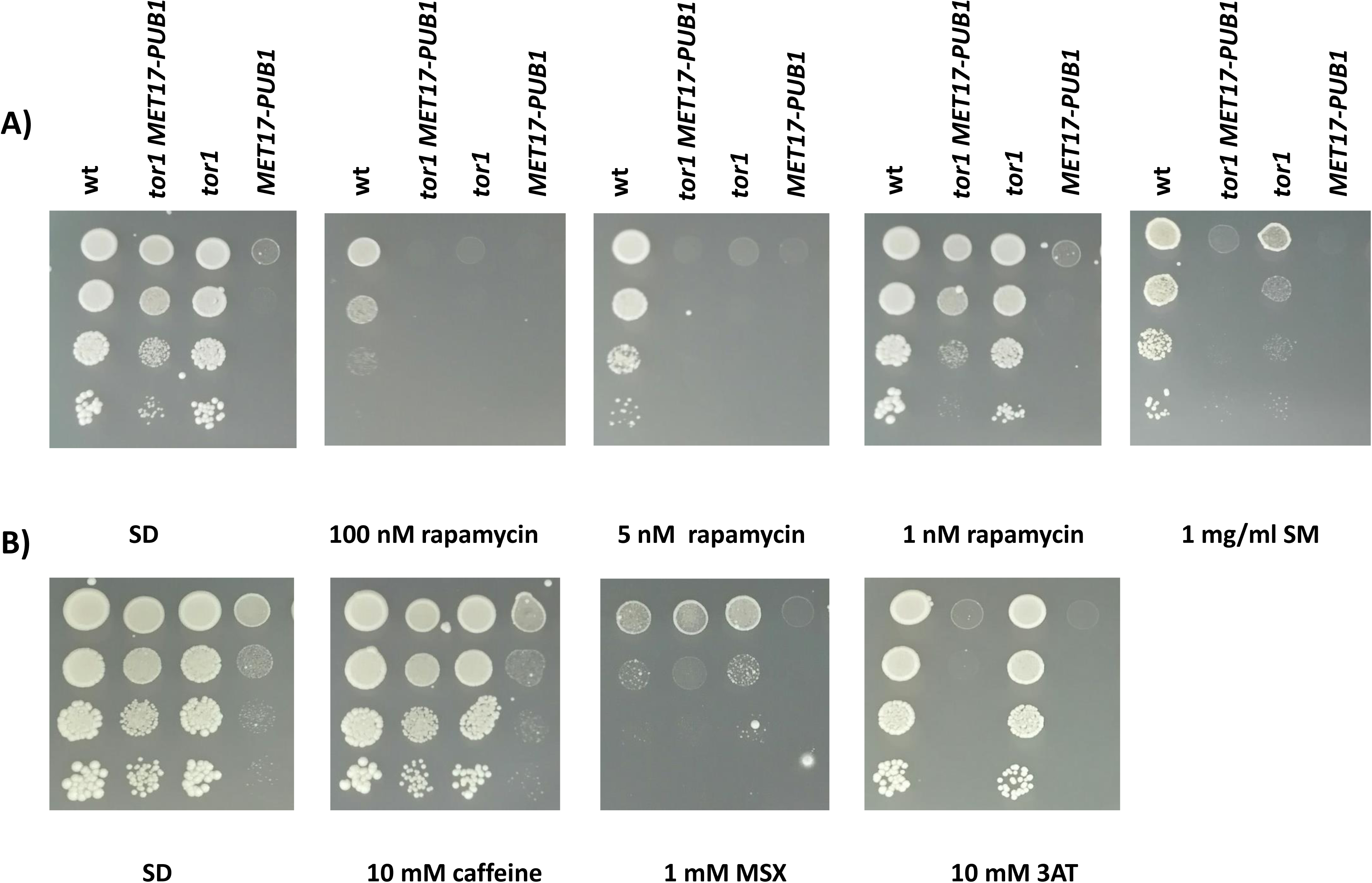
*PUB1* overexpression increases the sensitivity of cells to inhibitors of nutrient sensing pathways. A) The spot analysis of the dilutions of the wild-type, *MET17-PUB1, tor1*Δ, and *MET17-PUB1 tor1*Δ strains on minimal medium plates containing different amounts of rapamycin and 1 mg/ml sulfometuron methyl. Β) Same as panel A) on the plates containing 10 mM caffeine, 1 mM methionine sulfoximine (MSX) and 10 mM aminotriazole (3AT).

Caffeine is another molecule that has been proposed to inhibit, among other cellular processes, TORC1 (27). However, neither *TOR1* deletion alone nor combined with *PUB1* overexpression caused increased sensitivity to caffeine (Figure 4B). Next several inhibitors of amino acid synthesis were tested. Methionine sulfoximine (MSX) inhibits glutamine synthetase, which causes internal glutamine depletion that, in turn, causes partial TORC1 inhibition (28). Once again, the tested mutants displayed no different resistance to this inhibitor. Two inhibitors of amino acid biosynthesis were then tested. 3-aminotriazole (3AT) inhibits histidine synthesis (29), activating the general amino acid control (GAAC) kinase Gcn2. Sulfometuron methyl (SM) blocks the synthesis of branched amino acids like valine and isoleucine, causing an impact on the transcriptome that resembles TORC1 inhibition and Gcn2 activation (30). In SM, *PUB1* overexpression increased the sensitivity to these compounds more than *TOR1* deletion did alone (Figure 4A). This effect became even more dramatic in the 3AT-containing plates. Gcn2 deletion is hypersensitive to these compounds, so Pub1 may play a role in the GAAC pathway that may contribute to impaired growth in minimal medium. However unlike *TOR1* deletion, mutating the *GCN2* kinase gene or its primary target transcription factor *GCN4* did not rescue the growth of the *MET17-PUB1* mutant (Supplementary Figure S3A). Neither did the combination of *MET17*-*PUB1* and the deletion of yeast AMPK kinase homologue Snf1, involved in the aggregation of TORC1 component Kog1, affect growth (Supplementary Figure S3B). So it seems the Pub1 effect on growth is channelled mainly through TORC1 independent on Kog1. Therefore, Pub1 plays a complex role in amino acid biosynthesis control that may be sensitive to different nutritional deficits and/or might differently modulate TORC1 specific outputs.

### Pub1 controls Rps6 phosphorylation levels

In order to further study the role of Pub1 in TORC1 activity, we analysed the effect of the deletion and overexpression of the *PUB1* gene on the phosphorylation level of TORC1 target ribosomal protein S6 (Rps6). Rps6 is phosphorylated by the Ypk3 kinase in a TORC1-dependent way and is quickly dephosphorylated when TORC1 is inhibited by rapamycin (31, 32). Cells were grown exponentially on synthetic complete SC medium and two TORC1 inhibiting conditions were tested: rapamycin itself and nitrogen starvation (Figure 5A). As expected, rapamycin addition significantly reduced the phosphorylation of Rps6 in the wild-type strain. The amount of phosphorylated Rps6 protein was higher in the *PUB1* deletion mutant, even in control samples without rapamycin, supporting that the mutation resulted in enhanced TORC1 activity or inhibited Rps6 dephosphorylation. The total amount of Rps6 was also slightly increased, which indicates that the higher amount of phosphorylated protein may also partly reflect the amount of the total protein, or that phosphorylation stabilised the protein. *PUB1* overexpression appeared to elicit the opposite effect by causing the complete dephosphorylation of the protein in the presence of rapamycin, suggesting that TORC1 was more sensitive to inhibition. Nitrogen starvation caused complete Rps6 dephosphorylation in all the tested strains, which indicates that alternative pathways exist and may control Rps6 dephosphorylation in a Pub1-independent way. In order to gain an increased understanding of Rps6 dephosphorylation dynamics over time, samples were taken from 5 minutes to 1 h following the addition of rapamycin (Figure 5B). In the wild type, there was no evidence for dephosphorylation following short incubation times (from 5 to 15 minutes), whereas after 30 minutes dephosphorylation was evident whereas after 1 h it was almost complete. In the *pub1*Δ mutant, phosphorylation remained mostly unaltered for 30 minutes and only became evident after 1 h. Once again, not only the magnitude, but also the dephosphorylation rate increased in the *PUB1* overexpressed mutant: reduced Rps6 phosphorylation became evident after only 15 minutes and was almost complete at 30 minutes. Therefore, Pub1 appears to play an inhibitory role in TORC1-dependent Rps6 phosphorylation.

**Figure 5.**
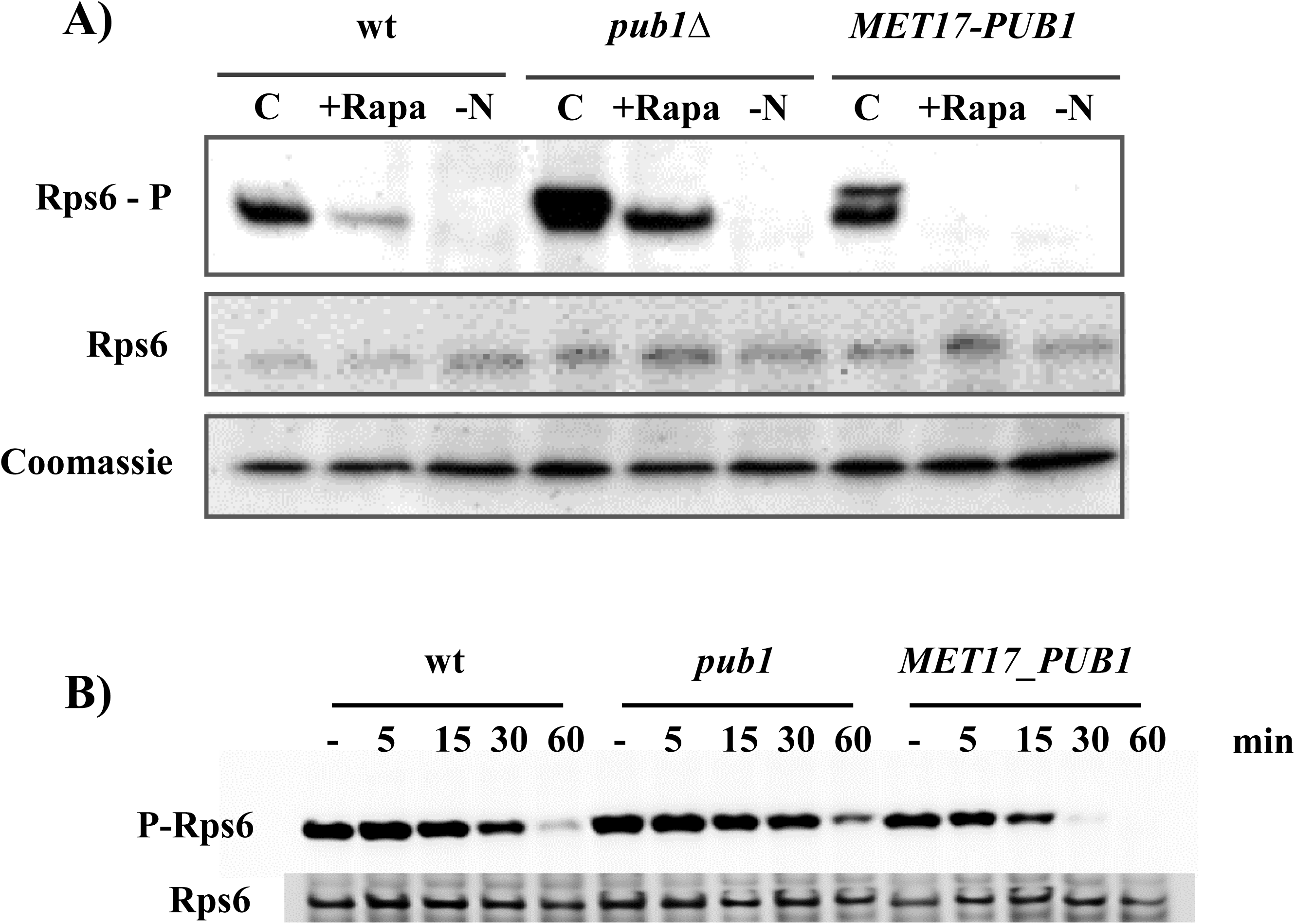
Pub1 inhibits Rps6 dephosphorylation by rapamycin. A) The Western blot analysis of the phosphorylation status of ribosomal protein Rps6 in the wild-type, *pub1*Δ and *MET17-PUB1* strains after incubation in 40 nM rapamycin for 30 minutes (+Rapa), or following nitrogen starvation for 1 h (-N). The control (C) indicates exponential growth in SC medium. A phosphorylated version of Rps6 was detected using the Phospho-S6 Ribosomal Protein (Ser235/236) antibody. Total Rps6 was used as the loading control. B) Rps6 dephosphorylation upon treatment with 40 nM rapamycin and sampling following 5, 15, 30 and 60 minutes.

### Pub1 overexpression triggers TORC1 aggregation

Pub1 is a protein with prion-like properties that is able to aggregate and is required for the formation of aggregates like SG. TORC1 is also sequestered transiently into granules after heat shock (8), and TORC1 component Kog1 aggregates in the so-called Kog1 bodies upon sugar starvation (33). Therefore, the aim was to investigate the role of Pub1 in potential TORC1 aggregation. To do so, we GFP-tagged the *KOG1* gene in the wild-type, *pub1*Δ mutant and *PUB1* overexpressing strains. *KOG1* is an essential gene, but the haploid strains expressing the fusion protein grew well, which indicated that the fusion was active. The effect of *PUB1* was analysed upon exponential growth in minimal SD medium when the protein was fully expressed (Figure 6A, Supplementary Figure S4A). Under this condition, Kog1-GFP neither aggregated in the wild-type strain nor in the *PUB1* deletion strain, as expected in a situation with no starvation. It was noteworthy, however, that multiple Kog1-aggregates were evident in the *MET17-PUB1* strain, even in the absence of stress. When cells enter stationary phase, aggregates are formed in the wild type strain, and their localization was similar to what was observed in the *MET17-PUB1* strain. However, such foci are absent in the *PUB1* deletion strain, indicating that Pub1 is required for their formation. Next we assayed whether Pub1 overexpression was also able to cause the aggregation of a Tor1-GFP fusion. According to the literature, the *TOR1*-GFP fusion leads to a non-functional protein (34) and, indeed, our fusion appeared to be hypersensitive to rapamycin (data not shown). Nevertheless, the Tor1-GFP protein accumulated in aggregates upon *PUB1* overexpression (Supplementary Figure S4B). Similarly to Kog1-GFP, foci were formed in the wild-type strain when cells entered the stationary phase. However, as with the Kog1-GFP fusion protein, foci were absent in the *pub1*Δ strain, which indicates that Pub1 is required to form these Tor1 aggregates. Glucose starvation is another stress condition known to induce the formation of Kog1 bodies (9). Therefore, we grew the aforementioned strains exponentially and shifted to the minimal medium with no carbon source for 2 h (Figure 6). The Kog1-GFP aggregates formed normally in the wild type and still formed when *PUB1* was overexpressed, but not in the *PUB1* deletion strain. These results indicate that Kog1-GFP foci that formed upon glucose starvation also required Pub1 to form.

**Figure 6.**
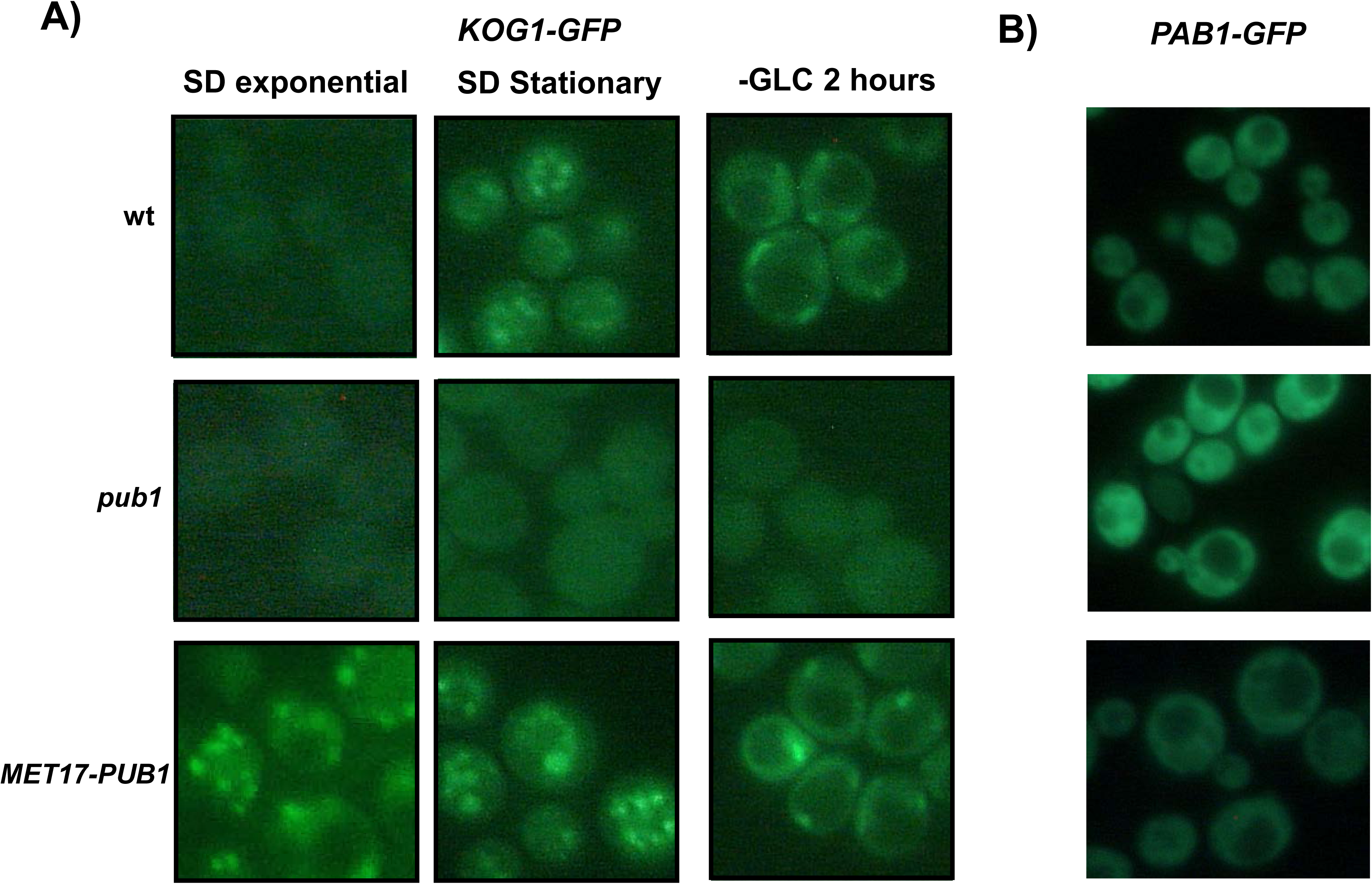
Pub1 is required for Kog1 aggregation. A) The wild-type, *pub1*Δ and *MET17-PUB1* strains expressing Kog1 tagged with GFP were observed under the microscope upon exponential growth in minimal medium SD in the stationary phase with the same medium and upon glucose starvation. B) The same strains, but expressing instead a GFP-tag on the *PAB1* gene, were observed during exponential growth in SD medium.

Another SG component, the poly(A) binding protein Pab1, was tagged with GFP in the same strains and was assayed under identical conditions (Figure 6B). In this case, fluorescence was always diffuse and did not form any aggregate. Therefore, the Kog1 aggregates induced by *PUB1* overexpression are not strictly equivalent to SGs. The aggregation of pyruvate kinase Cdc19 has been linked to SG formation and has been shown to inhibit TORC1 (35). Therefore, Cdc19 was tagged under the same conditions, but there were no signs of aggregation (data not shown). This suggests that the effect of Pub1 on TORC1 activity is not mediated through Cdc19-mediated. Overall, Pub1 overexpression impacted on Kog1 distribution, which might explain its other phenotypes as Kog1 aggregation has been shown to down-regulate TORC1 activity (9).

## Discussion

In this manuscript, the relationship between two apparently unrelated factors, e.g. the RNA binding protein Pub1 (with multiple functions in different cellular processes) and the TOR kinase of complex TORC1, was analysed. Tor1 can be replaced partially with Tor2 kinase in the formation of the TORC1 complex, but not to a full extent as mutant *tor1*Δ is hypersensitive to TORC1 inhibitor rapamycin (Figure 2). This allows the epistatic analysis of *PUB1* and *TOR1* deletions. Although *pub1*Δ is slightly more tolerant to rapamycin, *tor1*Δ*pub1*Δ is highly sensitive, which indicates that TORC1 acts downstream of Pub1 with regard to this particular phenotype. The fact that *TOR1* deletion also blocks the deleterious effects of Pub1 overexpression on both growth and longevity (Figures 3 and 4) reinforces the idea that Pub1 influences these processes through the TORC1 complex. The negative effect of high Pub1 levels on growth is more obvious with nitrogen starvation and reduced TORC1 activity, and suggests that the negative regulation of TORC1 function by Pub1. In contrast, Pub1 does not interact with other nutrient sensing pathways, such as the GAAC through the Gcn2 kinase or glucose repression through AMPK kinase Snf1 (Supplementary Figure S3). However, Gcn2 and TORC1 mutually regulate each other (36–38), and at this stage it is too early to rule out any interactions under some conditions or indirect effects.

Pub1 is an RNA-binding protein that associates with the translation machinery and contains prion-like motifs that allow it to self-aggregate. Pub1 binds to 368 transcripts (39), but TORC1 components are not included among them. Although Pub1 is a well-known translational repressor, its overexpression does not cause a drop in global polysome amounts, which implies that its function does not affect global translational initiation. However, the ability to regulate the formation of TORC1 foci under certain environmental conditions may be the key function of Pub1 in this case. SG formation depends on Pub1 being present under many environmental conditions (15). It has been described that TORC1 is sequestered transiently in the SGs that form under heat shock (8), and the aggregation of Kog1 reduces TORC1 signalling (9). We have shown that Pub1 overexpression triggers the formation of the cytoplasmic foci containing TORC1 components. Interestingly under the same conditions, the proteins associated with SG, such as Pab1 and Cdc19, do not aggregate. Hence the nature of these foci differs from SG under this particular condition. The Kog1-bodies described under glucose starvation tend to be present as a single focus per cell (9). The fact that we observed many aggregates when Pub1 was overexpressed indicates that they may constitute a different structure. There is no genetic interaction with the *SNF1* gene (Supplementary Figure S3), the kinase required to form Kog1-bodies, supporting this idea. Further analyses are necessary to determine if such foci are SGs but, in any case, the aggregation and/or mislocalisation of TORC1 may suffice to diminish its signalling power, delaying growth and enhancing rapamycin sensitivity. The fact that the composition of the SG forming in response to different stress conditions can be distinct has previously been observed following azide and glucose starvation (15). A similar TORC1 inhibitory function has recently been described for another SG component, Pbp1 (12). This is based on the ability of Pbp1 to self-associate into assemblies that are required for TORC1 inhibition under respiratory conditions. Therefore, a variety of proteins capable of self-aggregating may regulate TORC1 under different environmental conditions forming different intracellular condensates.

Pub1 impacts TORC1-controlled processes, such as autophagy (Figure 1) and Rps6 phosphorylation (Figure 5), but not with all stimuli. When Rps6 is dephosphorylated or Pgk1-GFP degradation by autophagy is triggered by rapamycin, Pub1 is clearly required. However upon nitrogen deprivation, both Rps6 dephosphorylation and autophagy activation occurs in a Pub1-independent manner. Pub1 may act on a subset of TORC1 complexes or near the subunit that interacts with rapamycin. The fact that several RBP behave differently argues against a model in which the SG structure is key to control TORC1. There is no evidence for the localisation of SG in the vicinity of the vacuolar EGO complex that is known to control TORC1. None of the published global phosphoproteomic analyses indicate Pub1 phosphorylation, although it is subjected to ubiquitinylation according to global studies, and its levels are regulated by the acetylation machinery (40). We cannot rule out the notion that TORC1 may play a role in Pub1 activity, but there is no evidence for TORC1 in SG formation. However when *PUB1* is overexpressed, the repression caused in global translation by rapamycin is partly suppressed, which suggests that some TORC1 functions may be channelled directly or indirectly through the same functions targeted by Pub1.

To conclude, our data add to recent studies that have suggested a key role of mRNA metabolism and ribonucleoprotein complexes in the regulation of nutrient signalling (41, 42), and also suggest a novel role of mRNA binding Pub1 in inhibiting TORC1 signalling.

## Materials and methods

### *S. cerevisiae* mutant strains construction

Mutant strains derive from prototrophic haploid strain C9 (43). Gene deletions were performed by the short end homology strategy using the recyclable selection marker *lox*P-*kan*MX-*lox*P contained in plasmid pUG6 (44). The antiobiotic resistance marker was excised whenever necessary by transforming with a plasmid containing recombinase *cre* and a cycloheximide resistance gene (45). *PUB1* overexpression was achieved by replacing its promoter with the *MET17* inducible promoter by using plasmid pkanMX-MET17p as previously described (19). The GFP tagging of *PGK1*, *KOG1* and *PAB1* involved the use of plasmid pFA6-GFP-KanMX6 (Longtine et al., 1998) as a template. All the yeast transformations were performed by the lithium acetate method (46).

### Yeast growth media and conditions

The standard medium for yeast growth was YPD (1% yeast extract, 2% bactopeptone, 2% glucose) (47), with 2% agar when the solid medium was required and geneticin G418 (at 20 μg/ml) to select for transformants. Minimal medium SD contained 0.17% yeast nitrogen base, 0.5% ammonium sulphate and 2% glucose (47). This medium was used to select the transformants with cycloheximide resistance by employing 2 μg/ml. The complete minimal medium SC used for the CLS was SD plus 0.2% of a drop-out mix with all the amino acids.

CLS assays were performed in SC medium (22). The cells grown overnight on YPD were used to inoculate SC media at an OD_600_ of 0.1. Aliquots were taken with time, diluted and plated. Colonies were counted and the percentage of survival was calculated by considering day 3 of growth to be 100% survival. The growth curves in the YPD medium were obtained by a Varioskan Lux plate reader with shaking and incubation a 30°C.

### Western blotting

The cells to be used for the autophagy and Rps6 phosphorylation assays were broken in a volume of glass beads in lysis buffer (Tris-HCl 0.1 M, pH 7.5, NaCl 0.5 M, MgCl_2_ 0.1 M, NP40 1% (v/v), PMSF 10 mM, protease inhibitors and phosphatase inhibitors) for three 20-second cycles in a FastPrep 24 shaker. Extracts were centrifuged at 4°C and full speed for 5 minutes to remove cell debris and the amount of protein in the supernatant was quantified by the Bradford method (Biorad Inc. Hercules, CA, USA). Extracts were mixed with the SDS-loading buffer and were subjected to electrophoresis in an Invitrogen mini-gel device. The gel was blotted onto PVDF membranes for Western blot analysis in a Novex semy dry blotter (Invitrogen, Carlsbad, CA, USA). Phospho-S6 Ribosomal Protein (Ser235/236) antibody (1:1000 dilution) was obtained from Cell Signalling Technology (Beverly, MA, USA). The anti-Rps6 antibody (1:1000 dilution) was acquired from Abcam (Cambridge, MA, USA). Anti-GFP (1:1000 dilution) was purchased from Santa Cruz Biotechnology (Santa Cruz, CA, USA). The secondary HRP-conjugated antibodies were purchased from Santa Cruz Biotechnology to be used at a dilution of 1:5000. The ECL Western blotting detection system (GE) was used following the manufactureŕs instructions.

### Polysomal fractionation

Cells were grown exponentially in SC media to an OD_600_ of 0.5-0.6. A portion of the culture (80 ml) was recovered and chilled for 5 min on ice in the presence of 0.1 mg/ml CHX. Cells were harvested by centrifugation at 6,000 × *g* for 4 min at 4°C and resuspended in lysis buffer (20 mM Tris-HCl, pH 8, 140 mM KCl, 5 mM MgCl_2_, 0.5 mM dithiothreitol, 1% Triton X-100, 0.1 mg/ml CHX, and 0.5 mg/ml heparin). After washing and resuspension in 700 μl of lysis buffer, a 0.3 ml volume of glass beads was added, and cells were disrupted 3 times in a Bio 101 FastPrep for 20 s at speed 5. Lysates were cleared by being centrifuged at 5,000 rpm for 5 min, after which the supernatant was recovered and was centrifuged at 8,000 rpm for 5 min. Finally, glycerol was added to the supernatant at a final 5% concentration before storing extracts at −80°C. The samples of 10–15 A_260_ units were loaded onto 10-50% sucrose gradients and were separated by ultracentrifugation for 2 h and 40 min at 35,000 rpm in a Beckman SW41 rotor operating at 4°C. Gradients were then fractionated using isotonic pumping of 60% sucrose from the bottom, which was followed by a recording of the polysomal profiles by online UV detection at 260 nm (Density Gradient Fractionation System; Teledyne Isco, Lincoln, NE. USA).

### Fluorescence microscopy

The GFP-labelled cells were directly observed in synthetic medium in different growth stages. Cells were visualised with an FITC filter under a Nikon Eclipse 90i fluorescence microscope.

## Funding

This work has been funded by grants PID2021-122370OB-I00 to EM and AA funded by MCIN/AEI/ 10.13039/501100011033 and by the European Union FEDER program, and the Swedish Research Council to MM. HO was a post-doctoral VALi+d fellow of the Generalitat Valenciana (APOSTD-2016-134).

## Author contributions

Planned experiments: HO, EM, MM, AA. Performed experiments: HO, AA. Analysed data: HO, EM, MM, AA. Contributed reagents or other essential material: EM, MM,

AA. Wrote the paper: AA.

## Conflict of interest

The authors declare no conflict of interest.

## Notes

### Competing Interest Statement

The authors have declared no competing interest.

