## Supplementary Figures 1-4 for "The RNA binding protein Pub1 inhibits TORC1 activity in *Saccharomyces cerevisiae*"

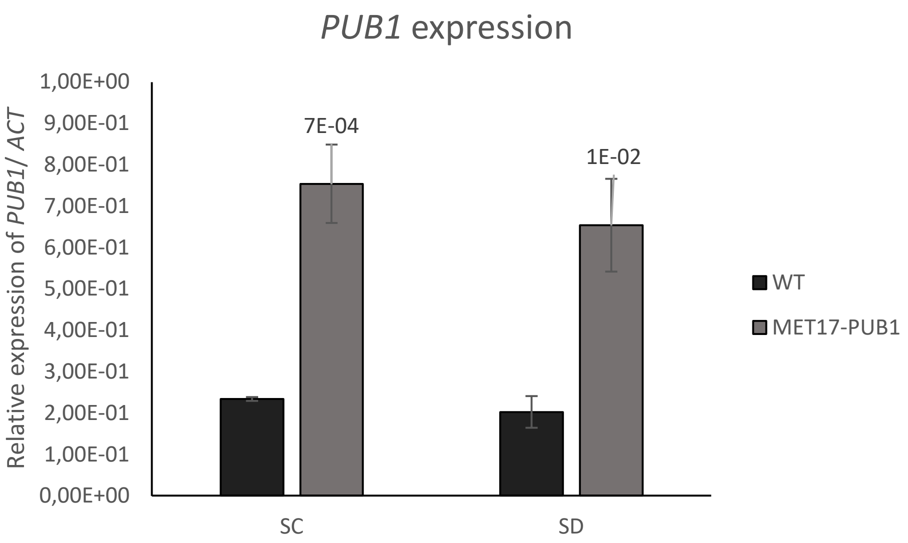


**Supplementary Figure S1**. qPCR analysis of PUB1 expression in the overexpressing strain *MET17-PUB1* vs the wild type C9 strain. Total RNA was amplified using the 5 x HOT FIREPol® EvaGreen® qPCR Mix Plus (Solis Biodyne) in the CFX Connect Real-Time System (BioRad).

### Control

### wt

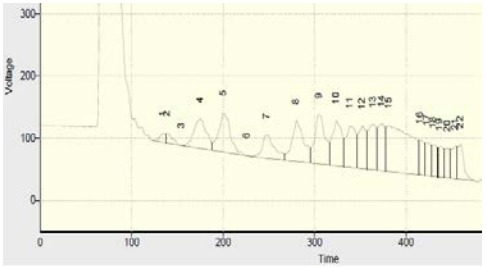


| | |
|---|---|
| Polysomes(%) | 85,50 |
| Polysomes/monosomes | 5,82 |

#### *tor1*

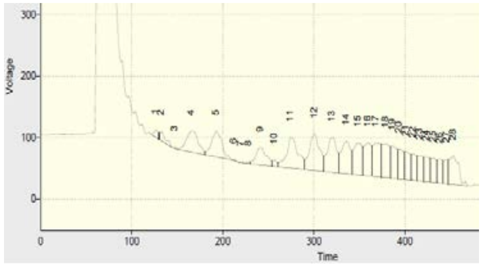


| | |
|---|---|
| Polysomes(%) | 84,90 |
| Polysomes/monosomes | 5,66 |

#### *pub1*

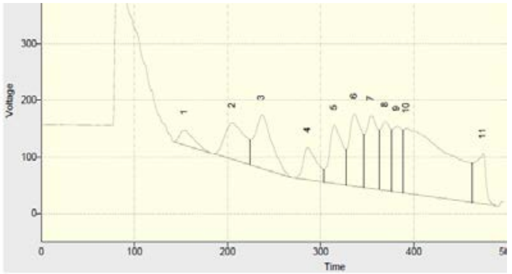


| | |
|---|---|
| Polysomes(%) | 90,70 |
| Polysomes/monosomes | 9,75 |

#### *MET17-PUB1*

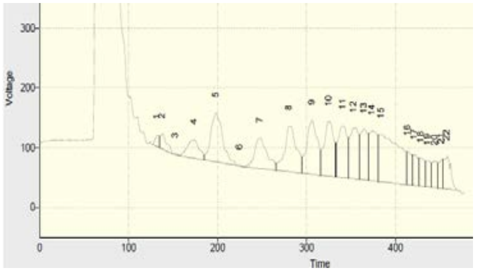


| | |
|---|---|
| Polysomes(%) | 85,80 |
| Polysomes/monosomes | 10,50 |

#### *MET17-PUB1 tor1*

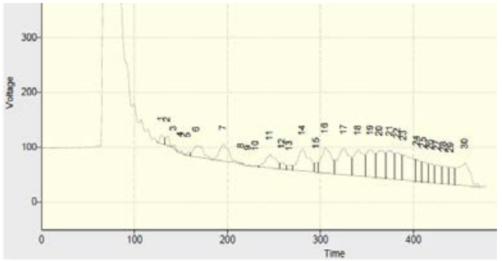


| | |
|---|---|
| Polysomes(%) | 87,70 |
| Polysomes/monosomes | 7,13 |

### +Rapamycin

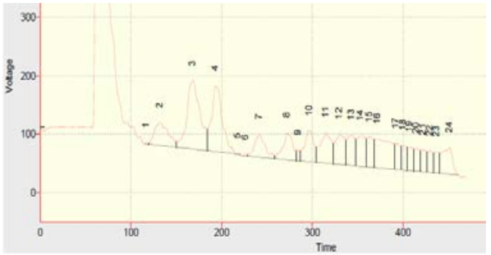


| | |
|---|---|
| Polysomes(%) | 66,40 |
| Polysomes/monosomes | 1,96 |

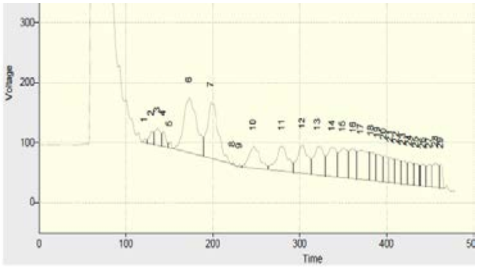


| | |
|---|---|
| Polysomes(%) | 69,90 |
| Polysomes/monosomes | 2,32 |

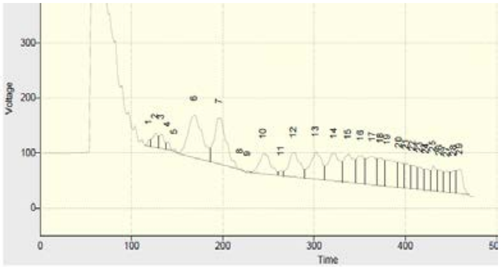


| | |
|---|---|
| Polysomes(%) | 72,90 |
| Polysomes/monosomes | 2,68 |

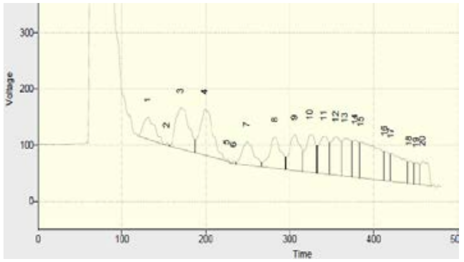


| | |
|---|---|
| Polysomes(%) | 75,20 |
| Polysomes/monosomes | 3,03 |

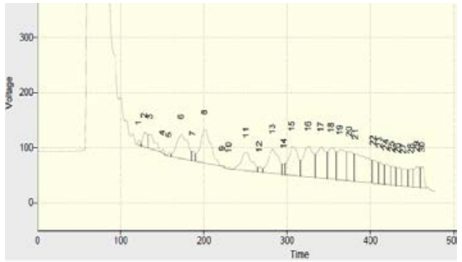


| | |
|---|---|
| Polysomes(%) | 79,60 |
| Polysomes/monosomes | 3,90 |

**Supplementary Figure S2.** Polysomal profiles of the indicated cells under exponential growth (control) or with 200 nM rapamycin for 15 minutes

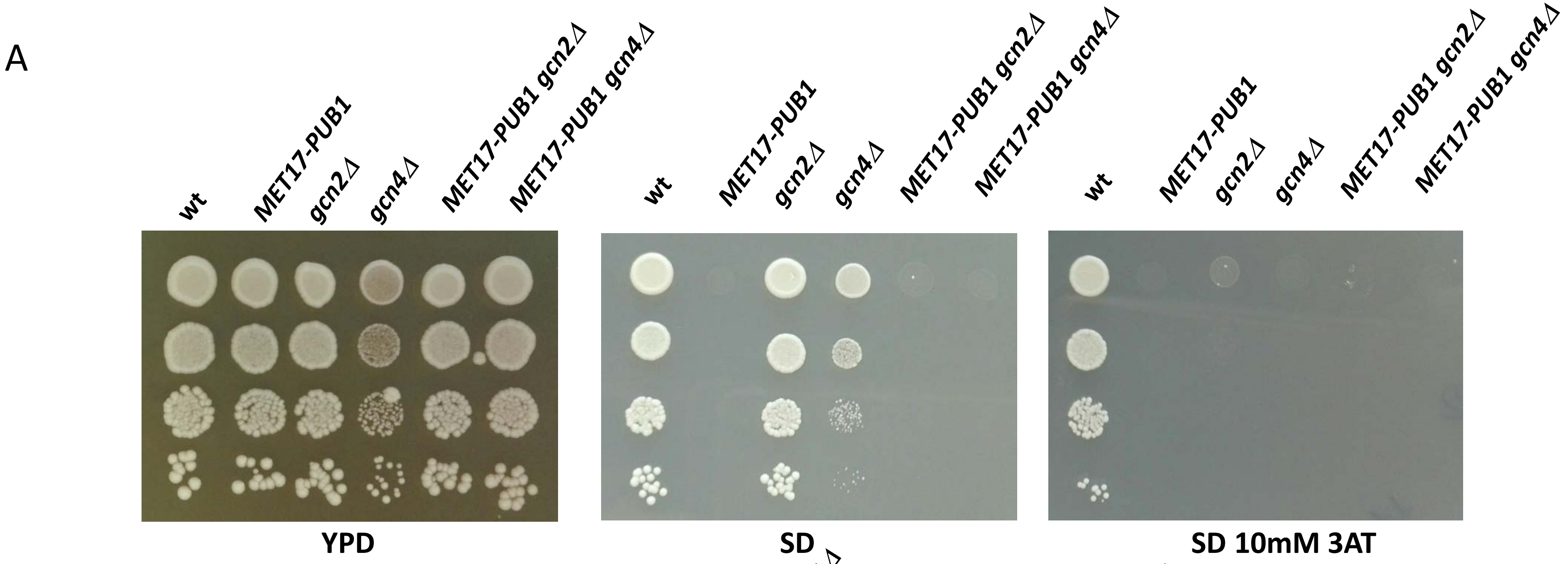


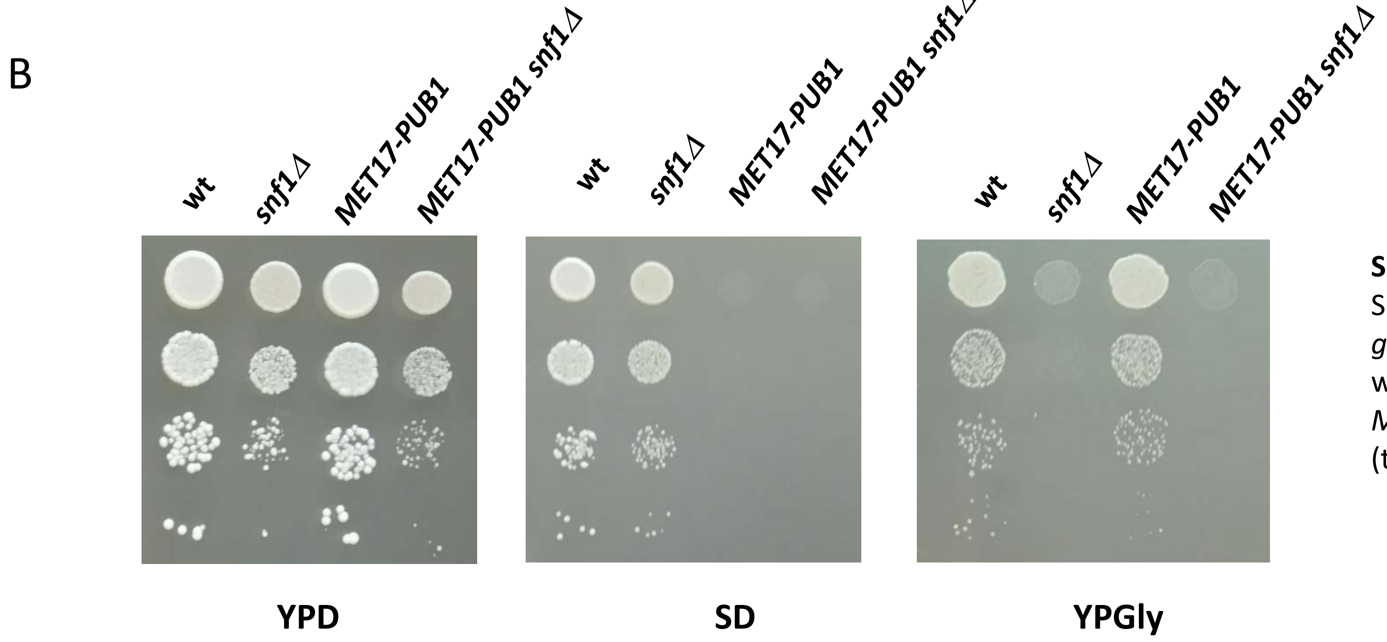


**Supplementary Figure S3**: PUB1 does not interact genetically with GAAC and Snf1 pathways. A) Spot analysis of dilutions of wild type, *MET17-PUB1*, *gcn2*Δ, *gcn4*Δ (and their double mutants) o rich medium YPD, on minimal médium SD with or withour 3-aminotriazol. B) Spot analysis of dilutions of wild type, *MET17-PUB1*, *snf1*Δ and *MET17-PUB1 snf1*Δ on YPD, SD and YPGlycerol plates (to test their ability to grow on non-fermentable carbon souces)

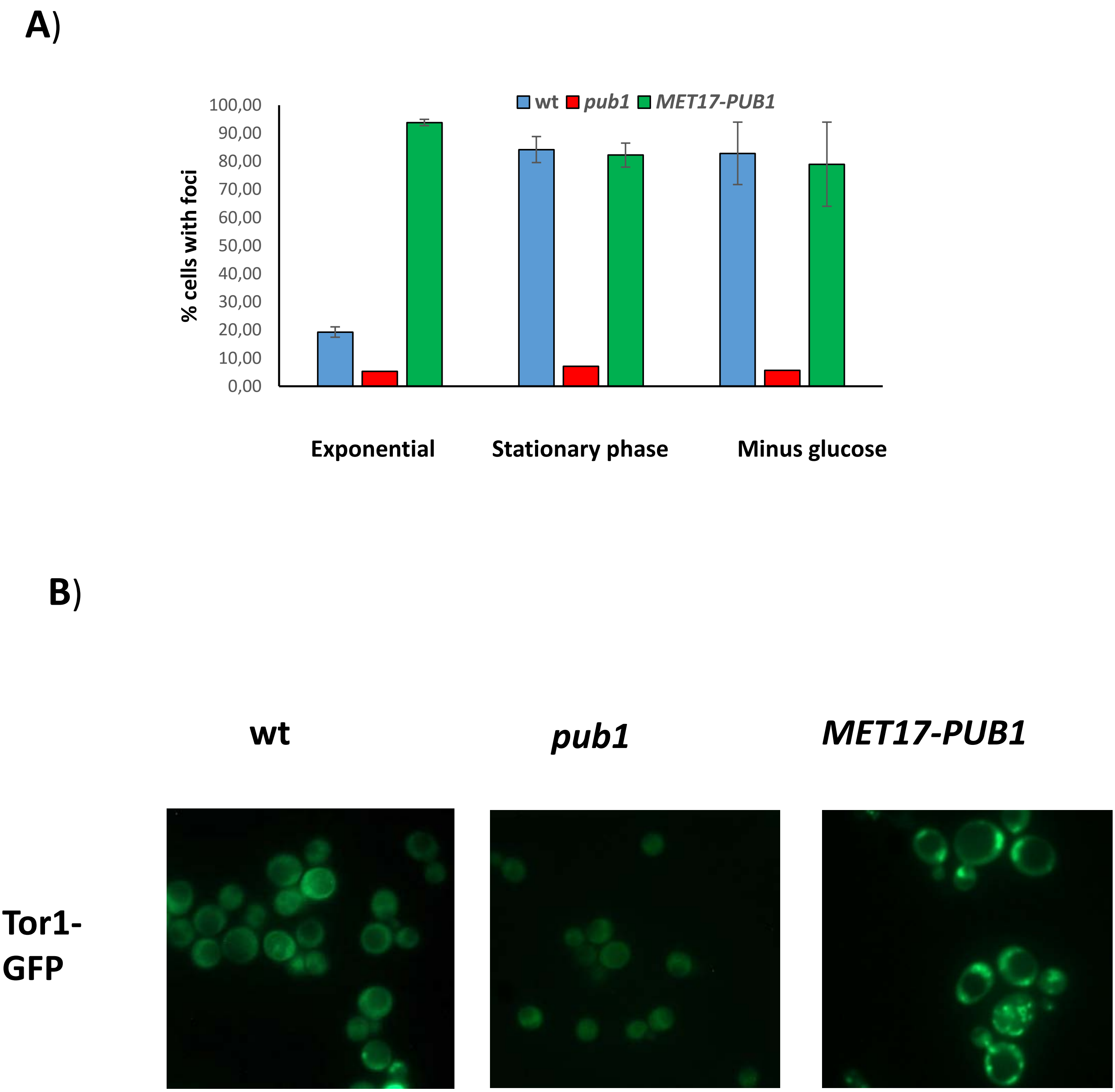


**Supplementary Figure S4.** Pub1 controls TORC1 localization. A) Cuantification of percentage of cells with foci form Figure 6A). Over 100 cells were counted for each mutant and condition.

B) TOR1-GFP visualization under exponential growth on minimal medium SD for the indicated strains
